# Spontaneous retinal waves generate long-range horizontal connectivity in visual cortex

**DOI:** 10.1101/2020.03.18.997015

**Authors:** Jinwoo Kim, Min Song, Se-Bum Paik

## Abstract

In the primary visual cortex (V1) of higher mammals, long-range horizontal connections (LHCs) are observed to develop, linking iso-orientation domains of cortical tuning. It is unknown how this feature-specific wiring of circuitry develops before eye opening. Here, we show that LHCs in V1 may originate from spatio-temporally structured feedforward activities generated from spontaneous retinal waves. Using model simulations based on the anatomy and observed activity patterns of the retina, we show that waves propagating in retinal mosaics can initialize the wiring of LHCs by co-activating neurons of similar tuning, whereas equivalent random activities cannot induce such organizations. Simulations showed that emerged LHCs can produce the patterned activities observed in V1, matching topography of the underlying orientation map. We also confirmed that the model can also reproduce orientation-specific microcircuits in salt-and-pepper organizations in rodents. Our results imply that early peripheral activities contribute significantly to cortical development of functional circuits.

**Highlights:**

- Developmental model of long-range horizontal connections (LHCs) in V1 is simulated
- Spontaneous retinal waves generate feature-specific wiring of LHCs in visual cortex
- Emerged LHCs induce orientation-matching patterns of spontaneous cortical activity
- Retinal waves induce orientation-specific microcircuits of visual cortex in rodents

**Significance statement:** Long-range horizontal connections (LHCs) in the primary visual cortex (V1) are observed to emerge before the onset of visual experience, selectively connecting iso-domains of orientation maps. However, it is unknown how such tuning-specific wirings develop before eye-opening. Here, we show that LHCs in V1 originate from the tuning-specific activation of cortical neurons by spontaneous retinal waves during early developmental stages. Our simulations of a visual cortex model show that feedforward activities from the retina initialize the spatial organization of activity patterns in V1, which induces visual feature-specific wirings of V1 neurons. Our model also explains the origin of cortical microcircuits observed in rodents, suggesting that the proposed developmental mechanism is applicable universally to circuits of various mammalian species.

## Introduction

In the primary visual cortex (V1) of higher mammals, neurons are observed to respond selectively to the orientation of visual stimuli, and their preferred orientations are organized into columnar orientation maps^1^ (**Fig. 1a**). In addition, iso-domains of the same orientation preference in the map are linked together by long-range horizontal connections (LHC)^2^ (**Fig. 1b**). The clustering of V1 by LHCs is observed prior to eye-opening^3,4^, suggesting that LHCs emerge prior to visual experience. Despite extensive studies on LHCs, it is still unknown what their functional role is^5,6^, and how they emerge before eye-opening.

**Figure 1.**
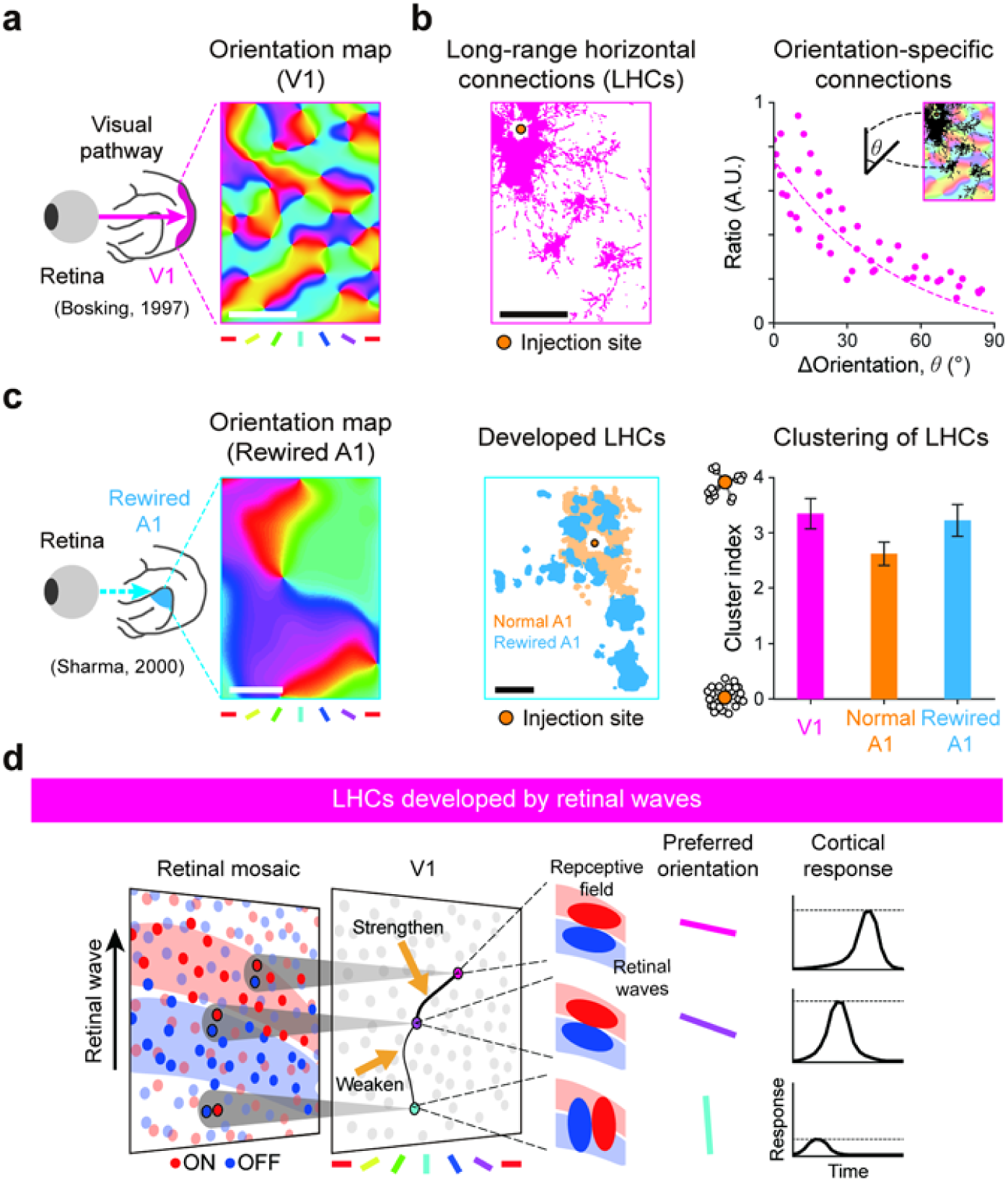
Tuning-specific horizontal connections in V1 can be developed by retinal waves. **a.** Orientation map in normal V1. Scale bar, 500 µm. **b.** Distribution of horizontal connections over the cortical space of (**a**), and its density in respect to orientation difference. Scale bar, 500 µm. **c.** (*Left*) Orientation map in rewired A1. (*Middle*) Distribution of horizontal connections in normal A1 and in rewired A1 showing that retinal afferent input induces the development of LHC-like long-range connections. Scale bar, 500 µm. (*Right*) Cluster index of horizontal connections. **d**. Illustration of the developmental model of orientation-specific connectivity by retinal waves. Following the statistical wiring model, local ON/OFF dipoles in an RGC mosaic are retinotopically wired to the V1, seeding cortical neuron anisotropic receptive fields and orientation tuning. The V1 contains a fully connected horizontal connection network, initialized with random synaptic strength. As a propagating retinal wave over the retinal mosaic co-activates cortical neurons

In previous studies, it has been suggested that the feedforward afferents from the retina may play a critical role in the development of cortical circuitry^7^. Sur et al. reported that rewiring retinal afferents to the primary auditory cortex (A1) in ferrets at early developmental stages results in development of orientation maps in A1^8^ (**Fig. 1c**). Notably, LHCs, not observed in normal A1, are observed to emerge in the rewired A1. These results suggest that retinal afferents initiate development of the cortical orientation tuning and LHCs during early developmental stages. This scenario was further supported by observations that orientation tuning of cortical neurons originates from the local ON and OFF feedforward afferents^9,10^. Moreover, the theoretical framework of the statistical wiring model indicates that the orientation tuning in V1 is constrained by the local structure of ON and OFF mosaics of retinal ganglion cells (RGCs)^11–13^ (**Fig. 1d**). These findings inspired our hypothesis that the structure of retinal afferents may induce feature-specific wirings of LHCs in V1.

Herein, we show that spontaneous retinal activity before eye-opening, which is spatio-temporally constrained by retinal mosaics circuitry, can selectively activate V1 neurons of similar orientation tuning and lead to developing LHCs via activity-dependent cortical plasticity. Our model is based on the observed data of spontaneous retinal activity^14,15^ and their coincidence with the development of the visual circuits^16,17^ (**Fig. 2**). It was reported that stage II retinal activity induces development of the retino-geniculate and geniculo-cortical pathways^18,19^, and then the stage III retinal activity that is observed until eye-opening^20,21^, coincides with the period of LHC development^22^. Considering that the retino-cortical projections are developed by the stage II retinal activity, we hypothesized that the stage III retinal activity is transmitted to the cortex and that this may play a crucial role in the development of LHCs.

**Figure 2.**
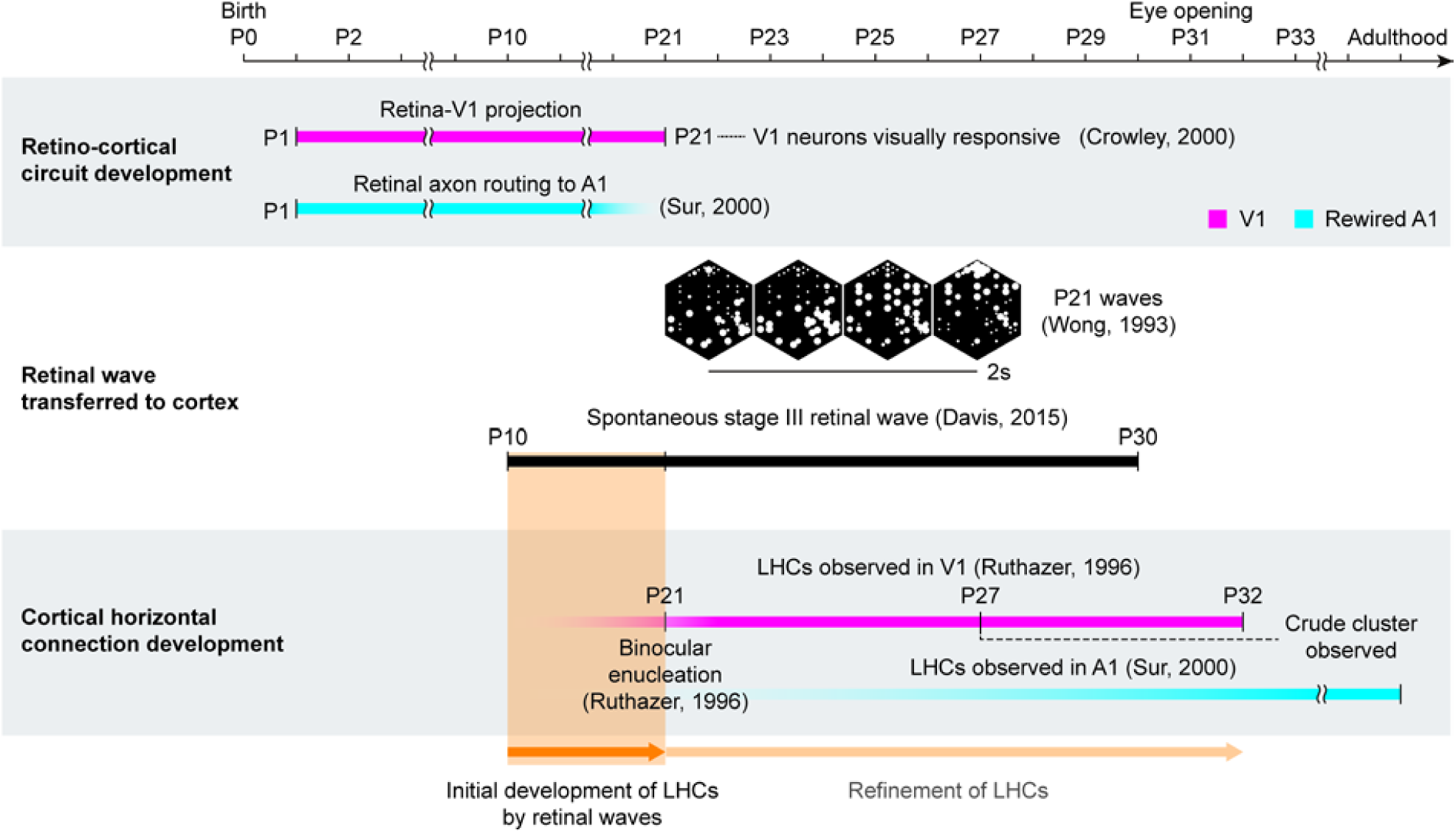
Development of long-range horizontal connections in V1 coincides with stage III retinal waves. A developmental timeline illustrates a coincidence between the emergence of LHCs and the spontaneous stage III retinal waves. A retino-cortical pathway is already developing at P10 when geniculo-cortical afferents reach layer 4^56^, and stage III retinal waves are observed from P10 until eye opening (P30)^22^. This suggests that V1 neurons can be activated by these waves from P10. Note that clusters of LHCs are observed, regardless of binocular enucleation at P21^3^, implying that orientation-specific LHCs can start emerging before then (Orange shaded area).

From the simulations based on the anatomy of retinal circuits, we demonstrate that temporally asynchronous retinal waves from ON and OFF RGCs can drive V1 neurons selectively by their orientation tuning, which drives tuning-specific wirings of neurons with activity-dependent plasticity^23,24^ of horizontal wirings in the cortex. We also show that the developed LHCs can induce patterned cortical activities, matching the topography of underlying orientation maps as observed in ferrets before eye-opening^25^. Finally, we demonstrate that LHCs can develop in the salt-and-pepper organization as observed in rodents^26^. These findings suggest that spontaneous retinal waves contribute significantly to organization of the functional architectures in the cortex during early developmental periods before sensory experience.

## Results

### Spontaneous retinal waves induced orientation-specific horizontal connections

We first implemented simulations of spontaneous retinal activity. We modeled retinal waves based on the propagation-readout model of the stage II waves by Butts et al.^27^ with necessary modifications for a model of the stage III waves. The actual retinal circuitry involving bipolar cells, diffusing glutamate and many more components, were simplified to a network containing ON/OFF RGC and cross-inhibitory amacrine cells (AC) (**Fig. 3a**). Based on the experimental observation that inhibitory transduction of amacrine cells induces temporal delay between the bursting activity of ON and OFF RGCs^21,28^, our retinal wave model (**Fig. 3b**) simulates spontaneous activity as follows: an ON wavefront is formed by local excitatory networks of ON RGCs. Then local ACs are excited by surrounding ON RGCs, which cross-inhibits neighboring OFF RGCs. The OFF RGCs are activated when local inhibition declines, forming an OFF wavefront. As a result, the simulated stage III wave at a given time appears as one of separate activities of local ON/OFF RGCs. Then, we simulated retinal waves using experimentally measured ON/OFF RGC cell body mosaics of a cat^29^ and a monkey^30^ (**Fig. 3c and d**, see also **Supplementary Video 1**). Because the RGC mosaic data lacked information about AC locations, we made a hexagonal AC lattice from the experimentally measured density.

**Figure 3.**
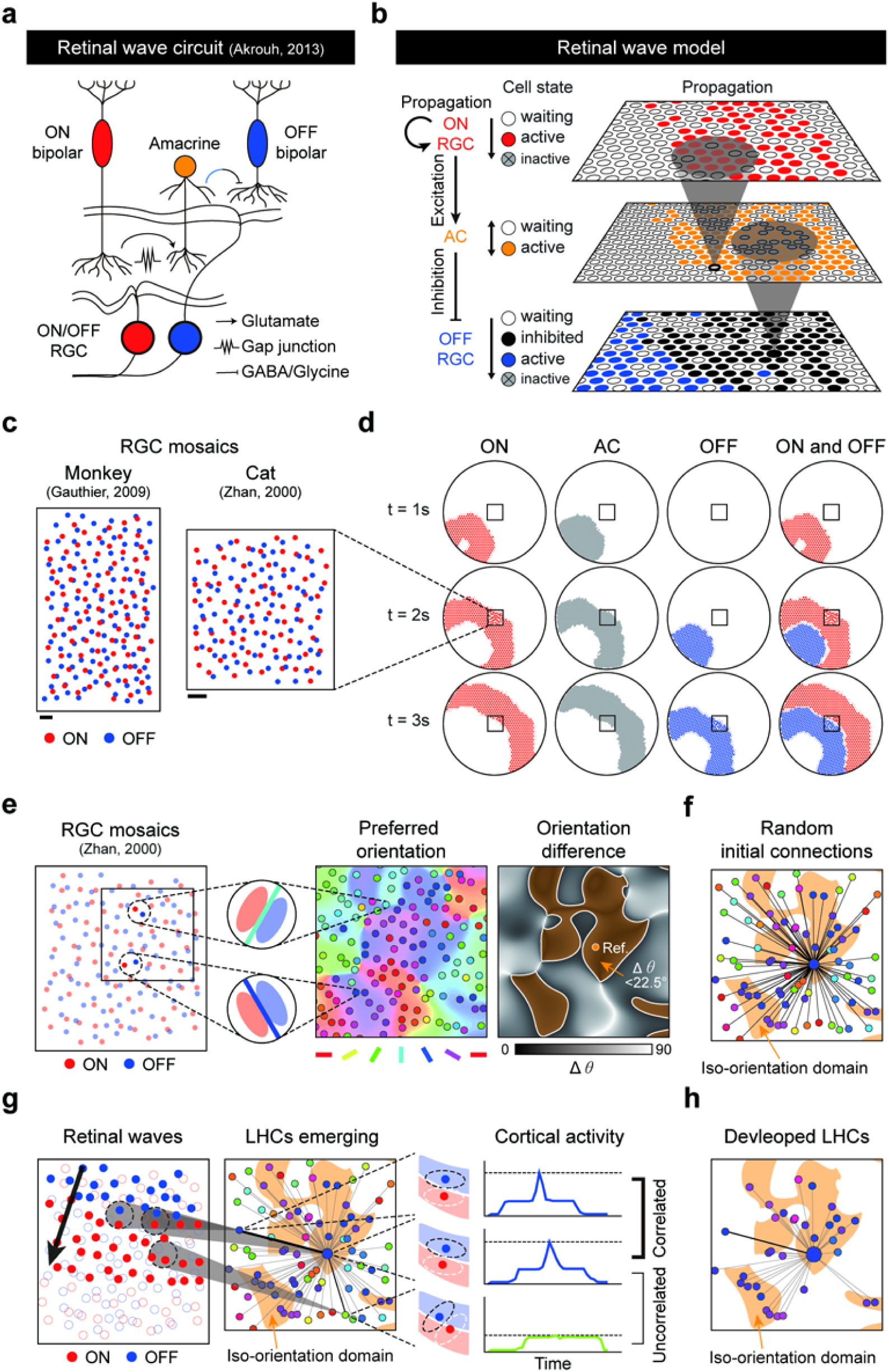
Orientation-specific horizontal connections develop by retinal waves. **a.** A simplified model of cross-inhibitory circuitry^57^ engaged during retinal waves. **b.** Simplified network a. model that simulates the cross-inhibitory behavior of amacrine cells and an illustration of waveform formation and propagation on overlaid hexagonal lattices of ON/OFF RGC and AC. **c.** The wave model was simulated using data RGC mosaics measured in cats^50^ and monkeys^30^. Scale bar, 100 μm. **d.** A simulated example of propagating retinal wave for a 3s period. Separation of ON/OFF activation regions is achieved by cross-inhibition by AC. **e.** (*Left and Middle*) The statistical wiring model in which orientation tuning is determined by retinal mosaics. (*Right*) Iso-orientation domains were indicated as yellow shaded area. **f.** Layout of the initial horizontal connections in V1. **g.** Simulation of the developmental model of orientation-specific connectivity by retinal waves. As propagating retinal waves provide a correlated activation of cortical neurons with aligned ON/OFF receptive fields, horizontal connections between neurons with the same orientation preference are selectively enforced by the Hebbian learning rule. **h.** Layout of the V1 horizontal connections developed by retina waves.

Next, following the statistical wiring model of the retino-cortical pathway^11,12,31^, we developed a model circuit of the neurons in V1. In this model, the receptive field of each V1 neuron is developed by retinotopic inputs from local ON/OFF RGC mosaics (**Fig. 3e**). The anisotropic alignment of ON and OFF receptive fields generates the orientation preference of each V1 neuron. In the current simulation, to mimic wirings of unrefined early cortical circuits, horizontal connections between V1 neurons were added with randomly initialized synaptic strengths (**Fig. 3f**, see Methods for details). Using this model, we simulated spontaneous generation of retinal waves and found that propagation of the ON and OFF retinal waves could provide a correlated activation of V1 neurons of similar orientation tuning (**Fig. 3g**). We confirmed that this correlated activation of V1 neurons was sufficient to strengthen their cortical wiring by a simple Hebbian plasticity implemented in horizontal connections (**Fig. 3h**).

After repeated propagations of spontaneously generated retinal waves in arbitrary directions, the initial random horizontal wirings turned to selective connections between neurons of similar orientation tuning (**Fig. 4a, b**). After training with approximately 500 waves of random directions, the statistics of LHCs developed by retinal waves showed significant bias of orientation-specificity as observed in ferrets^3^, whereas such biases are not observed in those developed by randomly permuted activities. (**Fig. 4c**, Cuzick’s test for trend; p = 0.3912, for n = 192,686 random initial synapses; *p = 4.94×10^−327^, for n = 173,437 synapses developed by retinal waves; p = 0.25, for n = 102,686 synapses developed by randomly permuted retinal activities). We observed that the cluster indices of developed LHCs in both cat and monkey models were comparable with those observed in Ruthzer et al. (1996)^3^, and were significantly higher than those developed by randomly permuted activities. (**Fig. 4d**, Developed (cat), two-tailed paired t-test, n = 50, *p = 2.43×10^−41^; Developed (monkey), n = 50, *p = 5.08×10^−6^).

**Figure 4.**
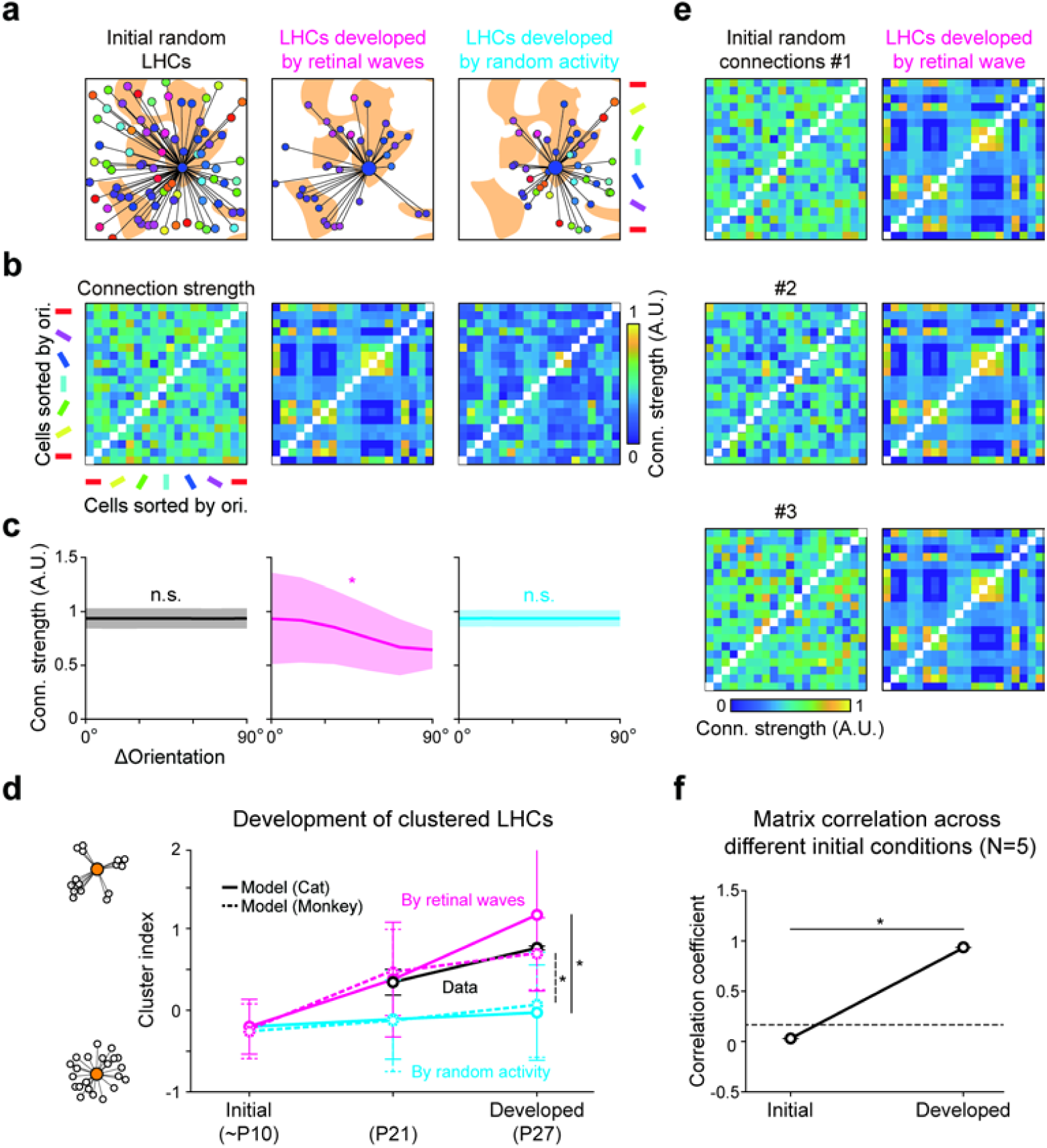
Organization of LHCs constrained by the structure of retinal afferents. **a.** Analysis of horizontal connection strengths in the initial random LHCs (*Left*), LHCs developed by retinal waves (*Middle*), and LHCs developed by random activity (*Right*). For each network, the locations of the top 10% of the strongest postsynaptic connections for a presynaptic location are shown over the orientation difference map. **b.** The connection strength (weight) for each pair of neurons was shown as a matrix. Cell indices were sorted by their preferred orientation. Shaded area indicates standard deviation. **c.** Orientation-specific LHCs developed by the model. Average connection strength was plotted as a function of orientation difference. **d.** Postsynaptic clustering in developing pre-EO ferret^3^ V1 and clustering in model V1 network developed from retinal mosaics in a cat and a monkey. Error bars indicate standard deviation. **e.** Repeated analysis for random initial connections. **f.** Correlation among connection strength matrices across different initial conditions (N = 5).

Next, to examine whether initial conditions of horizontal connectivity affect the developed structure of the orientation-specific LHCs, we repeated the simulation with different initial random horizontal networks (N = 5). As a result, we found that the horizontal connections develop in similar forms regardless of initial conditions before development (**Fig. 4e**). To analyze this result quantitatively, we estimated the correlation among the connection strength matrix of initial and networks developed across different initial conditions (**Fig. 4f**). We confirmed that correlation among the connectivity matrices of developed networks was significant (two-tailed paired t-test, n = 10, *p = 7.04×10^−25^). This result suggests that the structure of orientation-specific LHCs in the cortex develop under constraint by the retinal structure, regardless of the initial condition of connectivity.

### Spontaneous cortical activity induced by orientation-specific horizontal connections

Next, we examined whether the developed circuits of orientation-specific horizontal connections could reproduce patterns characteristic of the spontaneous cortical activity observed in early developmental periods. It has been reported that clustered cortical activity, topographically correlated with the underlying orientation map, is observed in developing visual cortex^32–34^ (**Fig. 5a-c**). A recent study on ferrets reported that the spontaneous cortical activity patterns before eye-opening predict the correlated organization of the orientation map in adults^33^ (**Fig. 5d**), suggesting that spontaneous activities in V1 may initialize the topographic maps in the cortex. However, this scenario could not explain clearly how spontaneously generated cortical activity organizes into systematic columnar patterns.

**Figure 5.**
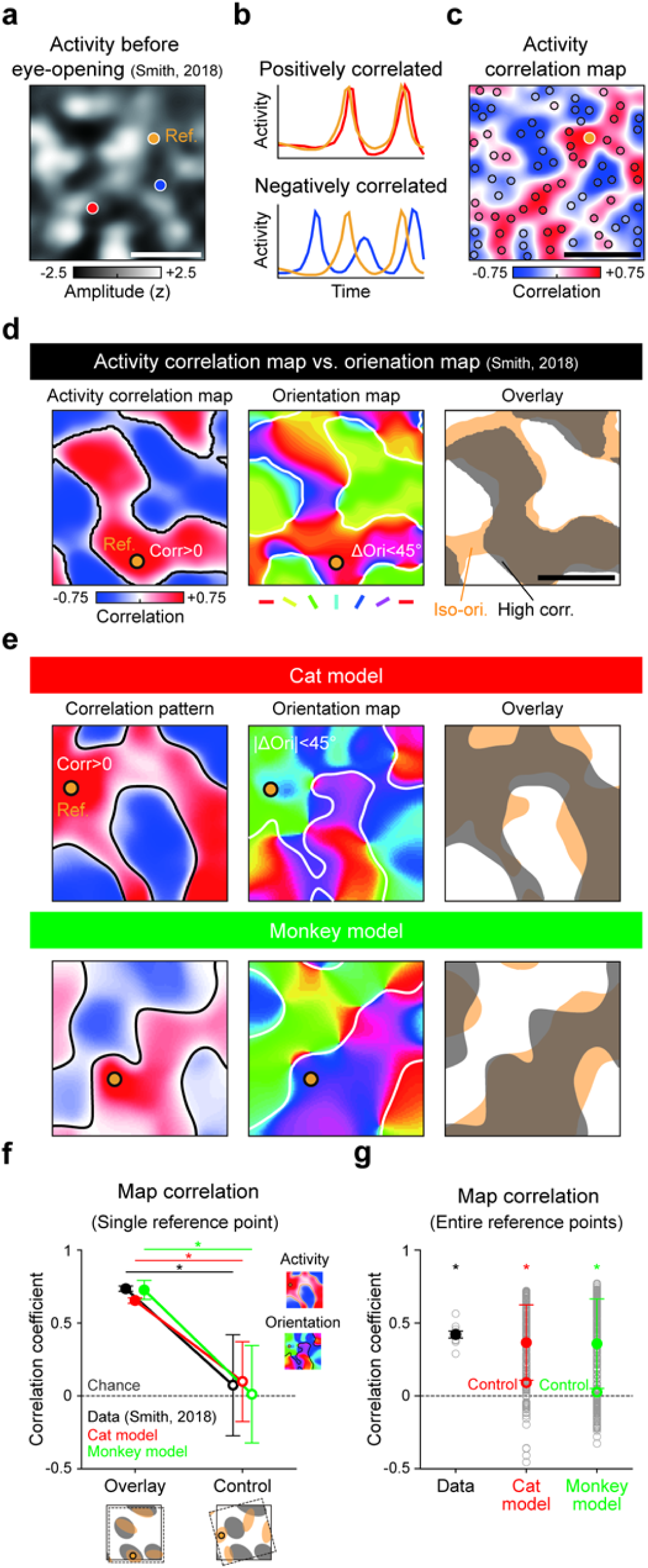
Orientation-specific connectivity developed by retinal waves can underlie correlated activity in early V1. **a.** Spontaneous events in early ferret V1 has an underlying correlation over cortical space. *Z*-scored images of spontaneous events observed in ferret V1 before eye-opening. Colored points indicate cortical locations of interest (orange: reference, red/blue: points to be compared with reference). **b.** Sample cortical activity correlations with respect to the reference point. Cortical activity at the red point has a higher correlation to the reference than the blue point does. **c.** Activity correlation map underlying early V1. Pearson correlation over the entire cortical space to the reference point is computed using complete activity images. **d.** Activity correlation map matched to orientation map in ferret V1. *Left*, Activity correlation map. The orange line denotes zero-correlation contour. *Middle*, Measured orientation map. The dark line denotes iso-orientation domain contours. *Right*, Alignment between positively correlated cortical regions and iso-orientation domains for a given reference point. **e.** Activity correlation map matched to orientation map in model V1 developed by retinal mosaic in cats^50^ and monkeys^30^. **f.** Correlation between activity correlation map and orientation similarity map in the data and model, as tested using Pearson correlation for a given reference point. Tested map pairs in (**e**). **g.** Correlation between activity correlation map and orientation similarity map in data and model tested for entire reference points (data: n=8, cat model: n=367, monkey model: n=318). Data adopted from Smith et al. 2018^33^.

Here, contrary to above scenario that spontaneously generated cortical activity pattern initially determines organization of the orientation map in V1, we suggest that spontaneous retinal activity determines the patterns of both orientation map and activity pattern in V1, by generating horizontal wirings that connect iso-domains of the underlying orientation map. In this simple scenario, topographies of spontaneous V1 activity and underlying orientation maps must be correlated. Further, once LHCs develop, silencing the feedforward activity in a retina or LGN^33^ cannot eliminate the correlated activity in V1, as observed in ferrets.

To validate this model, we removed all feedforward drive from the retina in our model and simulated activities by randomly driving the V1 network with developed LHCs. Following the analysis in a previous study^33^, we selected reference points at arbitrary locations in V1 and computed the Pearson coefficient of correlation for spontaneous activities between the reference and other locations across cortical space (**Fig. 5e**, left, see also **Supplementary Video 2, 3**). We observed strong matching between the activity correlation map and underlying orientation map, even though the V1 circuit does not receive inputs from the feedforward pathway. As observed in ferrets^33^, correlation between the activity correlation map and orientation map was significantly higher than in the controls where two maps were randomly rotated (**Fig. 5f**, two-tailed paired t-test with randomly aligned controls; data maps: n = 100, *p = 1.1×10^−35^, cat model: n = 100, *p = 2.23×10^−308^; monkey model: n = 100, *p = 1.90×10^−240^). We also repeated this for randomly chosen reference points and confirmed the statistical significance of the correlation (**Fig. 5g**, Mann-Whitney U-test; data maps: n = 8, p = 0.02, cat model: n = 367, *p = 1.17×10^−41^; monkey model: n = 318, *p = 4.94×10^−47^). These results suggest that the observed correlation between spontaneous V1 activity and orientation maps can readily be explained by our model.

### Development of orientation-specific horizontal connections without a periodic map

So far, we have shown that our model provides an explanation for how LHCs develop spontaneously in V1 of higher mammals with columnar orientation maps. Next, we show that our model further explains how orientation-specific LHCs also develop in rodents V1 with salt-and-pepper organizations^35^ (**Fig. 6a**). It is notable that the key assumption of our model is that retinal waves co-activate V1 neurons of similar tuning to develop LHCs between them, and that this mechanism works regardless of the spatial organization of orientation preference in V1 (**Fig. 6b**).

**Figure 6.**
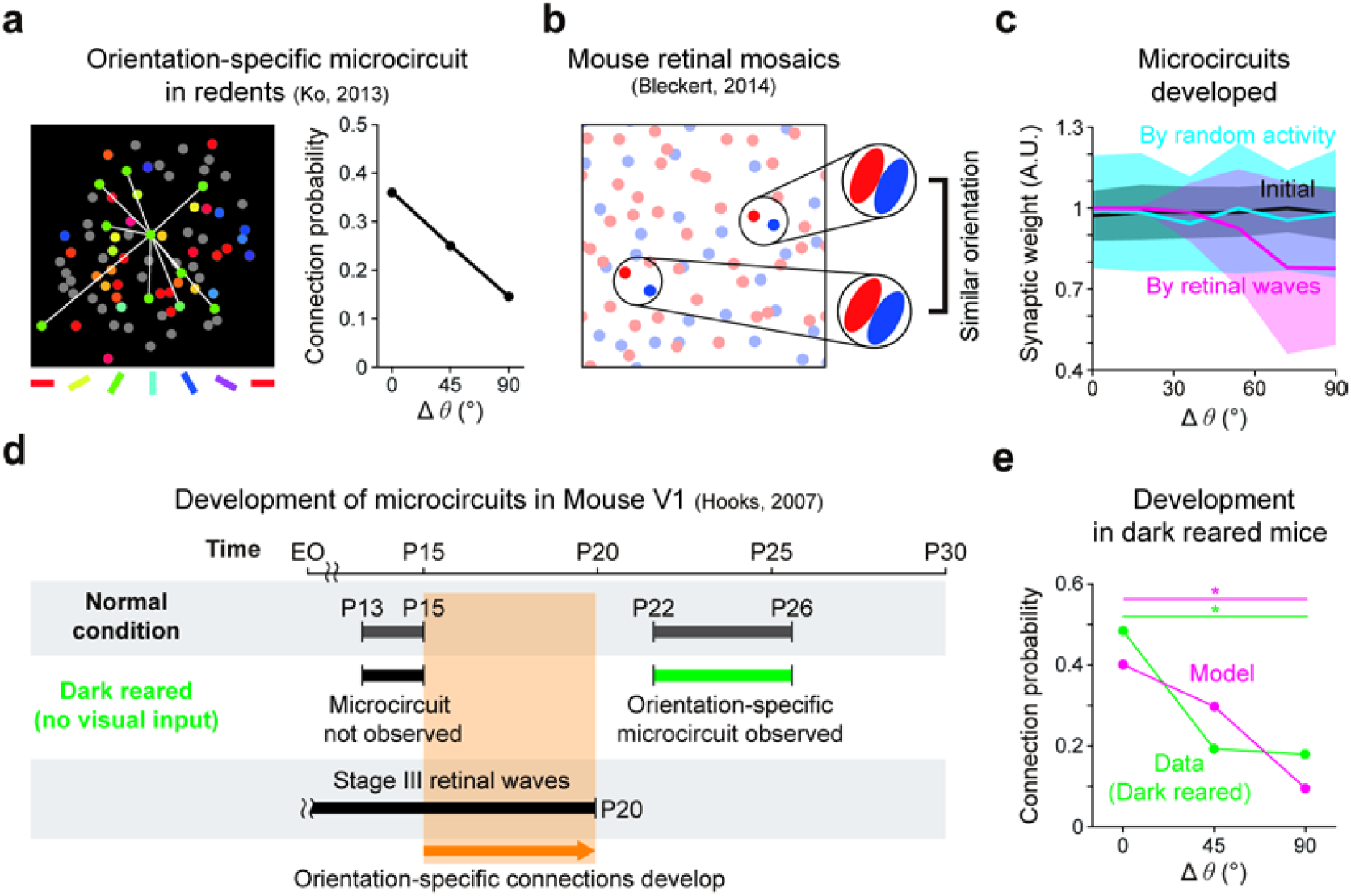
Retinal waves can drive the emergence of orientation-specific cortical connections in rodent V1. **a.** Orientation specific microcircuits observed in mouse V1 of salt-and-pepper organization of orientation tuning^45^. **b.** Emergence of orientation-specific horizontal connections by retinal waves in salt-and-pepper organization simulated with mouse retinal mosaics^36^. The model predicts that orientation-specific connections develop, regardless of spatial distribution of orientation preference in V1. **c.** Orientation-specific microcircuits developed by a simple orientation-correlated activation model. Shaded area indicates standard deviation. **d.** Developmental timeline of mouse V1 horizontal microcircuits. Orientation-specific horizontal microcircuits emerge when the retinal wave is present and the retino-cortical pathway has developed. **e.** Orientation specific connection developed in model mouse V1 and data^37^. In both model and data, the orientation specific microcircuit is observed even if no visual experience is given during development. Data adapted from Ko et al. (2014)^37^.

To validate our prediction in salt-and-pepper type organizations of V1, we implemented a model V1 circuit with salt-and-pepper organization. Using mouse retinal-mosaic data^36^ and the same developmental model for cats and monkeys, we confirmed that cortical neurons of similar orientation tuning tend to fire in correlated patterns. As a result, similar to the V1 model with a periodic orientation map, we found that LHCs with significant orientation-specificity developed (**Fig. 6c**, Cuzick’s test for trend; initial: n = 970, p = 0.15; developed by retinal waves: n = 551, *p < 2.23×10^−308^; developed by random activity: n = 970, p = 0.09). This result demonstrates that orientation-specific LHCs can also emerge from the correlated activity induced by retinal waves, even in salt-and-pepper type organizations^26^, as observed in layer 2/3 horizontal microcircuits of rodent V1^35^. Notably, observations that orientation-specific microcircuits appear to develop even in dark rearing conditions with no visual experience may also support our model. Our model predicts that these microcircuits can develop in dark rearing conditions, because spontaneous retinal waves can contribute under this condition^37,38^ (**Fig. 6d** and **e**, Cochran-Armitage test; Model: n = 551, p = 9.87×10^−13^; Data (Dark reared)^37^: n = 6, p = 0.028). These results imply that our model can provide a universal principle for the developmental mechanism of LHCs in both higher mammals and rodents.

## Discussion

Results from a number of studies have suggested that retinal waves may play an important role in the development of early visual circuits such as retino-geniculate pathways, geniculate receptive fields, and geniculo-cortical projections^22,39,40^. However, before now, it was not known whether retinal waves could also contribute to the early organization of functional circuits in V1. In the current study, we demonstrated that spontaneous retinal waves can drive the development of orientation-specific intracortical circuits in V1.

Previous studies have suggested that early V1 activity is driven by endogenous connectivity inside the cortex and plays a causal role in the development of horizontal connections. It was proposed that local circuits in V1 may generate patterned activities spontaneously, which serve as a scaffold for orientation columns to develop^33,41,42^. Furthermore, it was reported that early orientation maps are refined, matching spontaneous activity patterns during development before eye-opening. However, this scenario lacks explanation of a mechanism in which such organized activity patterns arise initially. Moreover, this model could not account for the strong and precise relationship between retinal and cortical receptive field structures reported recently^9,43^. Thus, the contribution of the endogenous cortical network alone does not explain how the tuning-specific horizontal connections develop and are correlated to orientation tuning in V1.

On the other hand, our model explains the emergence of correlated activity patterns in V1, the topographic matching to orientation map, and how these processes are carried out before the onset of any visual experience. Our result proposes a simple but powerful model of the developmental mechanism underlying the origin of spontaneous activity patterns in V1 and its correlation to orientation tuning maps, to complete the scenario.

Observations that support our developmental model were also reported in previous studies. Durack et al. (1996)^44^ found that initial clustering of LHCs in ferret V1 coincides with, but does not precede, the development of orientation preference^44^. This implies that the development of LHCs may “reflect”, rather than “seed” the structure of orientation maps. In addition, orientation-specific microcircuits of V1 in mice begin appear to emerge after development of retino-cortical projections and orientation tuning of V1 neurons^45^. These results also suggest that, after feedforward pathway has been developed to induce cortical orientation tuning, retinal waves drive the development of orientation-specific horizontal connections.

The previous study reported that clustered horizontal connections are observed even after binocular enucleation^3^. However, this result cannot invalidate the role of the patterned retinal activity for cortical development of LHCs. It should be noted that the enucleation of the retinae was performed late in development (P21), after early orientation tuning and related spatial organization were already established in V1. At that time, orientation-specific LHCs are expected to exist already, and might be refined by cortical activities^8^. Furthermore, another study reported a counterexample that the clustering of LHCs was not observed in cats with earlier binocular deprivation in the developmental stage ^7^. These results are readily explained by our model: Once orientation tuning and LHCs in V1 are established by early retinal afferents (probably early stage III), they develop without further contribution from the feedforward retinal activity in later stages.

Consistent with previous observations, our model proposes that stage II and III retinal waves may have distinct roles in development of LHCs. That is, the stage II retinal activity first induces development of the retino-geniculate and geniculo-cortical pathways^18,19^; then the stage III retinal activity drives the development of LHCs^22^ until eye-opening^20,21^. Another recent study on the stage II and III retinal activities in mice also provides supporting evidence for this model^46^. In this paper, it was reported that the global correlation between cortical activity across the cortical surface and the pre-synaptic retinal activity is significantly higher during stage II than during stage III. These results are consistent with the scenario that our model predicts: (1) Retinal activity during the stage II period develops retinocortical pathways with a retinotopic organization and activates all the cortical neurons. This generates a global correlation of cortical activity. (2) Then, asynchronous retinal waves during stage III selectively activate cortical neurons of similar orientation tuning by propagating in the direction of the wave. This leads to relatively lower global correlation than in the previous stage. Altogether, these results imply that stage II and III retinal activities stimulate the cortical neurons in a distinct way and contribute differently to the development of LHCs.

There exist conflicting results on whether the ON/OFF asynchrony of stage III retinal waves is significant in higher mammals. Wong et al. (1996)^47^ and Lee et al. (2002)^48^ reported that there was no noticeable temporal delay between ON and OFF RGC activity in ferret stage III waves, whereas Liets et al. (2003)^20^ reported the existence of clear asynchrony up to approximately 1 s, similar to the ON/OFF asynchrony in mouse. In the case of ferrets, it is possible that asynchrony of activity between RGC types might originate from a different mechanism, such as dendrite morphology or other neuronal dynamics, which eventually works similar to ON/OFF asynchrony in mice. Future studies are needed to observe detailed dynamics for retinal waves to drive cortical organization in each species of various structures in feedforward projections.

Our model explains how visual cortical networks develop before experience, but experimental results suggest that sensory experience is essential for the maintenance and maturation of cortical network structures. Immediately after eye-opening, an explosive increase in the synaptic density of cortical layer 2/3 (including LHCs) is observed. Ferrets dark-reared at this developmental stage appear to have much weaker orientation tuning, and their LHCs clusters do not form properly^49^. Similar observations were reported in rodent V1 as well; although orientation-specific cortical microcircuits are observed to develop in dark-rearing environments, they require visual experience for appropriate pruning and maturation later^37^. These observations suggest that early feedforward projections are able to guide organization of the functional circuits in the cortex initially, but also that visual experience is required to drive further development of circuits for complete visual function in adult animals.

In conclusion, our results suggest that the structure of retinal mosaics and the spontaneous wave activity from them can induce early tuning maps and the featured tuning-specific LHC circuits in V1. Our model provides further understanding of how functional architectures in the cortex can originate from the spatial organization of the periphery, without sensory inputs during early developmental periods.

## Methods

### Model simulations

Model simulations were designed based on the statistical wiring model from the retina to V1 pathway^11,12,31^, and performed the following: (1) Generation of spontaneous retinal waves, (2) Generation of orientation tuning of V1 neurons from statistical projections of RGC mosaics, (3) Development of horizontal circuits in V1 by retinal waves, and (4) Generation of spontaneous cortical activity from the developed V1 circuitry. Details of the algorithms, analysis methods, and parameters used in the simulations are presented in the following sections. All model simulations and data analyses were implemented and performed using MATLAB R2018a.

#### Retinal ganglion cell mosaics

For the simulation of our retina-V1 model, we used ON/OFF-type RGC mosaics data from mammals. Simulations shown in the main results are based on cell body mosaics of the cat^50^ (**Fig. 3-5**) and mouse^36^ (**Fig. 6**). We also provided simulation results based on monkey receptive field mosaics^30^. Considering that differences in cortical tuning organization, modeling, and simulation schemes differ between higher mammals (cat, monkey) and rodents (mouse); from here, we describe the framework for simulating higher mammalian visual cortex and later describe specifications for the rodent cortex model.

For an RGC mosaic, we denote positional vectors of ON/OFF-type cells as

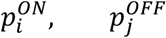

where *i* and *j* denote the cell index. We also measured the density of OFF RGC, and from that, we defined a representative spacing *d*_*OFF*_ so that an ideal hexagonal RGC lattice with spacing *d*_*OFF*_ would have the same cell density as the data OFF mosaic. We computed *d*_*ON*_ in the same manner.

#### Extending a data mosaic

To simulate spontaneous retinal waves before eye-opening, we modeled the retinal circuitry as a coupled network between ON RGC, OFF RGC, and amacrine cells (AC)^21^. Because the data mosaic represents a limited region of the retina (several hundred micrometers) and lacks AC positional information, we first augmented the data RGC mosaic with surround RGC padding and an AC lattice. Note that added cells were only used to aid wave propagation, and we did not consider them in further simulations.

First, to provide a retinal region sufficient for the wave to propagate, we padded the data RGC mosaic with synthesized hexagonal lattices of ON and OFF RGCs, each with spacing *d*_*ON*_ and *d*_*OFF*_. Note that with this spacing, a synthesized lattice has the same cell density as the data mosaic. We gave the padded lattice a circular boundary of radius 3000 µm (limited by GPU memory) to reduce potential propagation direction bias.

Then, a hexagonal AC lattice of (ON density + OFF density) was synthesized and overlaid on the augmented RGC mosaic. The resulting extended mosaic is an overlap of large ON/OFF/AC lattices with circular boundary, and we denote positional vectors of each type of cell as

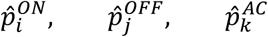

where *i, j*, and *k* denote the cell index. When referring to an extended mosaic, we use hat notation 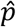 instead of the term *p* used for a data mosaic.

#### Network model generating retinal waves

In previous studies, it was reported that at least five cell types (including cone bipolar cells, amacrine cells (AC), and RGC) and several different transduction mechanisms, contribute to retinal waves. Based on the circuit mechanism reported by Akrouh et al. (2013)^21^ and the stage II wave model proposed by Butts et al. (1999)^27^, we designed a simplified retinal circuitry model involving ON/OFF/AC cells, where three types of local connectivity drove retinal wave propagation: ON→ON excitatory, ON→AC excitatory, and AC→OFF inhibitory^21,27^. Below we provide a conceptual description of our model network.

In our model, wave propagation happens within the ON mosaic layer, and AC/OFF layers serve as readout. We define the connectivity and activation rules among the layers as follows: (1) ON RGCs are reciprocally coupled with nearby active ON RGCs within the dendritic interaction radius *R*_*ON*_, and becomes active for 1 s when the summed input passes some threshold. (2) ACs receive input from nearby ON cells within *R*_*ON*_ and become active when the input passes a threshold. (3) Then, ACs provide inhibitory input to OFF RGCs within the dendritic radius *R*_*AC*_. An OFF RGC becomes inhibited with AC input and becomes active for 1 s upon reduction of such inhibitory input. The described connectivity is modeled as follows:

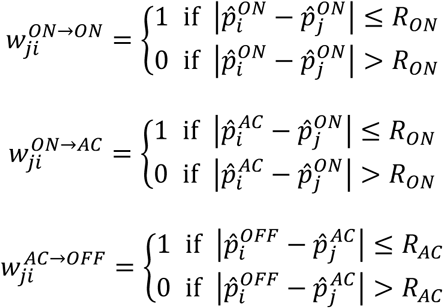

We chose the dendritic radius of ON RGC and AC considering the size of the cell dendritic arbors experimentally measured by Akrouh et al. (2013)^21^, but with some flexibility regarding the effect of diffuse neurotransmitters. Parameter details are provided in Table 1.

**Table 1.**
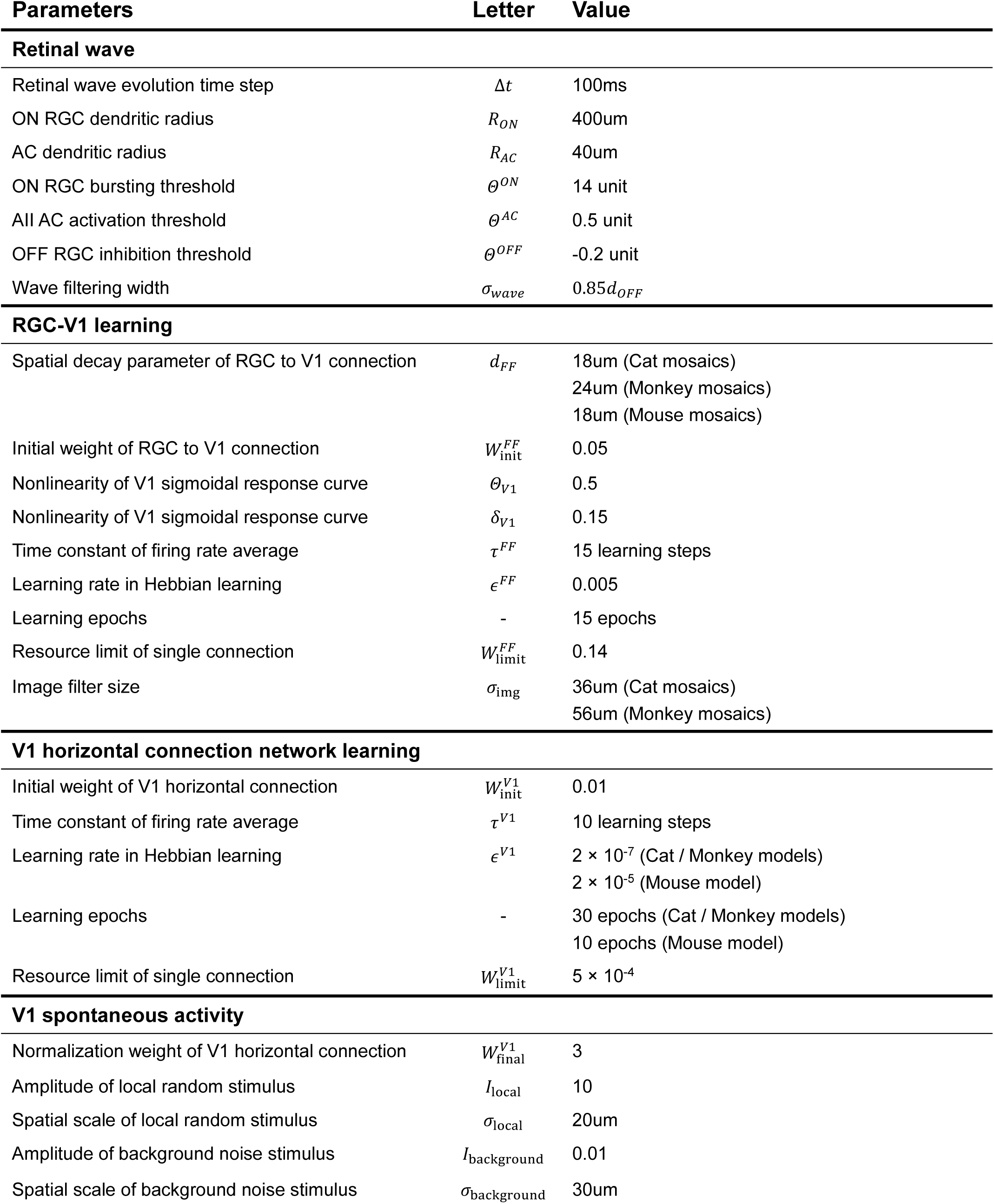
Parameters used in model simulation

#### Propagation mechanism of retinal waves

We designed our model based on a cellular automaton, in which each cell (at a specific time) is assigned a discrete cell state, and the cell states at time *t* + Δ*t* are determined by updating the cell states at time *t* according to a set of cell type-specific rules. We describe the cell type-specific states, input rules, and state update rules of our model below^27^.

The state of an ON RGC *S*^*ON*^(*t*) at a given time *t* can be *waiting, active*, or *inactive*. If an ON RGC is in a *waiting* state, it can switch to an *active* state at the next timestep by receiving input exceeding the threshold Θ^*ON*^ from nearby coupled ON RGCs in an *active* state (**Fig. 3b**). For realistic simulation, the amount of *η*_*c*_, which is the input that an *active* cell provides to other coupled cells, is drawn from a normal distribution of mean “1” and standard deviation Δ*C* = 0.2. After an ON RGC becomes *active*, it remains *active* for *T*_*a*_ = 1 *s*, after which it becomes *inactive* for the rest of the simulation.

The state of an AC *S*^*AC*^(*t*) can be either *waiting* or *active* at a given time. When a *waiting* AC receives input exceeding threshold Θ^*AC*^ from connected ON RGCs, it becomes *active* in the next time step. Different from RGCs, an AC switches back to *waiting* state whenever it receives input that does not exceed Θ^*AC*^, thereby retaining the ability to become active again.

The state of an OFF RGC *S*^*OFF*^(*t*) can be *waiting, inhibited, active*, or *inactive* at a given time. When a *waiting* OFF RGC receives inhibitory (negative sign) input not exceeding threshold Θ^*OFF*^ from connected ACs, it becomes *inhibited* in the next time step. When inhibitory input declines and allows the threshold to be exceeded, the OFF RGC state rebounds to *active*, which it maintains for *T*_*a*_ = 1 *s* before going *inactive* for the rest of the simulation. The parameter details are listed in Table 1.

#### Wave initiation, control, and postprocessing

For waveform and dynamics control, we initially assigned *waiting* state to only a portion (80%) of randomly selected ON RGCs and assigned *inactive* state to the remainder (20%)^51^. Then, we initiated a wave at *t* = 0 near the boundary of the extended mosaic by assigning *active* state to *waiting* ON RGCs within a circular region of radius *R*_*init*_ = 400 µ*m*. Afterward, the wave was allowed to propagate freely, while indices of *active* ON/OFF RGCs were recorded over time. For each mosaic, we simulated a set of 500-1000 retinal waves, each with different initial conditions and wave initiation positions.

Because the cells initially assigned *inactive* status cannot participate in the wave until the end (leaving “holes” in the propagating waves), we conducted a postprocessing process that ensured that all RGCs inside a propagating wave were densely *active*. At each time step, we spatially smoothed *active* state profiles of ON and OFF RGC layers using a 2D Gaussian filter of width *σ*_*wave*_ = 0.85*d*_*OFF*_. Then, we normalized the resulting activation values so that the maximum activation of each RGC layer was always “1” over time.

Finally, to ensure a uniform direction distribution, we classified the waves into 12 directional categories and sampled an equal number of waves from each category to construct a final, direction-unbiased wave dataset. From that, we imported random waves when needed.

Apart from patterned waves, we constructed a separate, permuted version of retinal waves as a control case for our horizontal network developmental model. The permutation was done simply by shuffling the RGC activation values over the RGC indices, thus preserving the overall activation level while removing correlated wave patterns.

#### Initialization of the RGC-V1 statistical wiring model

At the initial time of development, we assumed that RGCs are locally wired to cortical space retinotopically^12,31^. To determine cortical sampling locations, we looked for every pair of data ON/OFF RGCs with a distance less than 1.5*d*_*OFF*_ and set their center locations as cortical sampling sites. Note that we did not consider padded RGCs in the extended mosaic.

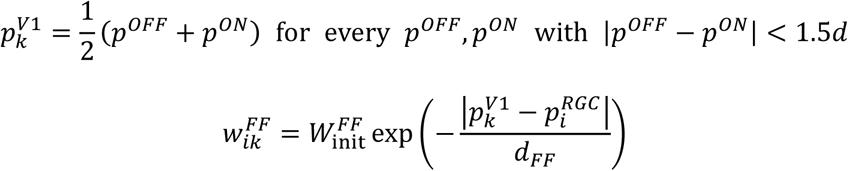

Here, 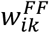 represents the feedforward connection weight from the *i*^*th*^ RGC to *k*^*th*^ cortical site, where 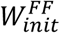 is the initial connection weight^52^. Parameter details are provided in Table 1.

#### Nonlinear response curve of V1 neurons

The response curve of V1 neurons was modeled as a nonlinear sigmoid kernel with parameters *δ*_*V*1_ and *Θ*_*V*1_.

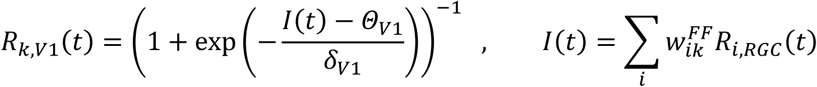

Here, *R*_*k,V*1_(*t*) is the response of the *k*^*th*^ cortical neuron at wave time *t*, where activation of *i*^*th*^ RGC by the retinal wave is given by *R*_*i,RGC*_ (*t*). Input to a cortical cell is solely determined by retinal feedforward input here, but horizontal input or direct cortical stimulus is allowed in later simulations. Parameter details are provided in Table 1.

#### V1 receptive field formation by retinal waves

After initialization with exponential pooling function, each ON/OFF subregion of the V1 “receptive field” is contributed by ∼1 RGC. To enlarge further the V1 receptive field before simulating the horizontal connection network, we followed the V1 receptive field developmental model of Song et al. (2018) to develop further the feedforward connections between the RGCs and V1 neurons^31^. For each feedforward connection, a weight update was done once per retinal wave, following the rule below.

For a given V1 neuron receiving input from a retinal wave, we sampled the peak response of the V1 neuron and related RGC activity levels at the same time. Then, we used profiles of the sampled responses from many retinal waves for an update of the covariance rule-based weights^53^, following the formulation below.

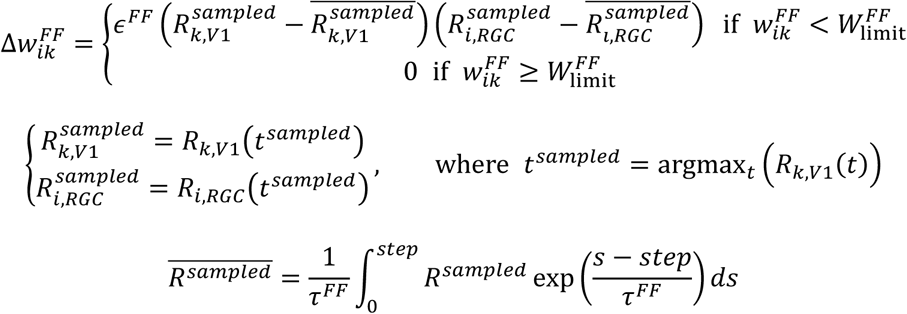

Here, we define the learning threshold 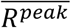 as the running average of the sampled responses of a cell during the learning steps. The term *τ*^*FF*^ represents how fast the threshold changes during the learning steps and the term *ϵ*^*FF*^, the learning rate, denotes how quickly the weight update is done. We assumed that the resource for a single connection is limited by 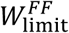.

We simulated the development of RGC-V1 feedforward connections using a post-processed retinal wave dataset, 15 epochs in total. At each epoch, we shuffled wave order and iterated over all waves one by one, applying the learning rule. All the parameter details are provided in Table 1.

#### Initialization of the V1 horizontal network model

After the RGC-V1 feedforward development was complete, we froze the feedforward connection weights and simulated development of the V1 horizontal connection network by retinal waves. Initially, we horizontally wired V1 cells with random weights.

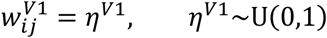

Here, 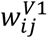 represents the synaptic connection weight from the *i*^*th*^ to *j*^*th*^ cortical site (*i* ≠ *j*), where *η*^*V*1^ is drawn from a random uniform distribution *U*(0,1). After setting the random connections, we normalized each V1 cell’s outgoing connection weight sum by 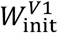 and finished the initialization step. Parameter details are provided in Table 1.

#### V1 horizontal network development by retinal waves

For each V1 horizontal connection, a weight update was done once per retinal wave. For a given retinal wave, responses of the V1 neurons over time were determined by the sum of feedforward input from RGCs and horizontal input from other V1 neurons.

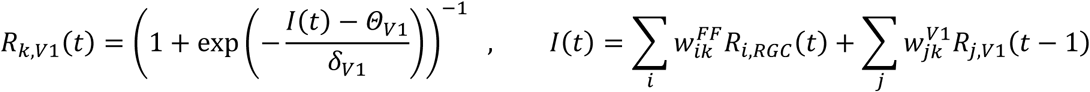

Then, peak responses of V1 neurons were sampled and used as a response profile for a covariance rule-based weight update of the V1 horizontal network.

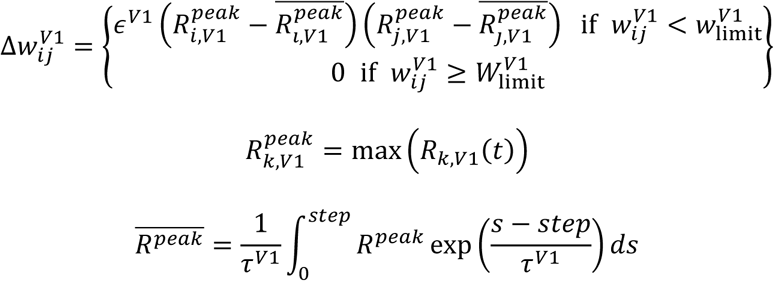

Here, we define the learning threshold 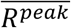 as the running average of peak responses of a V1 neuron over learning steps. Here, *τ*^*V*1^ represents how fast the threshold changes during the learning steps and *ϵ*^*V*1^, the learning rate, denotes how quickly the weight update is done. We assumed that resource for a single connection is limited by 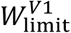.

We simulated development of the horizontal connection network using a post-processed retinal wave dataset (permuted dataset as control case), 30 epochs in total; at each epoch, we shuffled the wave order and iterated all the waves one by one, applying the learning rule. All the parameter details are provided in Table 1.

#### V1 horizontal network-driven spontaneous activity

After horizontal connection development was complete, we froze the horizontal connection weight and simulated spontaneous patterned activity induced by horizontal connectivity. We normalized each V1 cell’s incoming connection weight sum to 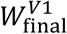 and modeled driving input *I*_stim_ for V1 network as a sum of local stimulus and global background noise, denoted by

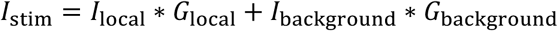

where *G*_*local*_ is the local dominant 2D Gaussian input of width of *σ*_local_ given at a random location in the cortical space with peak value “1”, and *G*_background_ is background random noise drawn at each cortical neuron from *U*(0,1), smoothed by 2D Gaussian filter of a width of *σ*_background_, and normalized to have a maximum value of “1”. The terms *I*_local_ and *I*_background_ denote the intensities of local stimulus and global background noise.

Then, in the absence of feedforward input (all *w*^*FF*^ = 0), we provided driving input *I*_stim_ to neurons in the V1 horizontal network and integrated their responses recurrently until the network activity diverged.

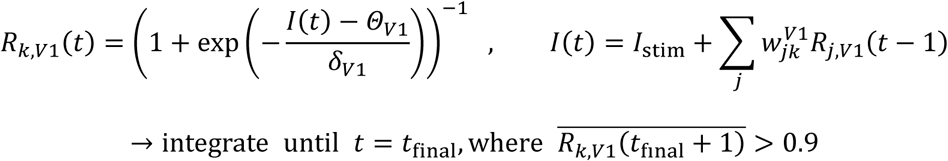

Then, we spatially filtered the cortical response profile of V1 neurons using a 2D Gaussian filter of width *σ*_img_ to obtain a response image *A*(*x*) where *x* denotes pixel position. After that, we normalized the image pixel intensities to be zero-centered and to have a standard deviation of “1”. Repeating the entire procedure, we modeled *N* = 200 spontaneous activity images, each indexed as *A*_*i*_ (*x*). Parameter details are provided in Table 1.

#### V1 spontaneous correlation patterns

Using the simulated spontaneous activity images *A*_*i*_(*x*), we computed spontaneous correlation patterns as the pairwise Pearson’s correlation between activity at a reference pixel *s* and activity at all other pixels *x*, given by

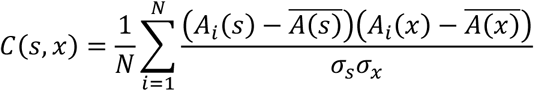

Here, the horizontal bar denotes averaging over all spontaneous activity images and *σ*_*x*_ denotes the standard deviation of activity over all images at location *x*^33^.

#### Measurement of the cortical orientation map

We calculated the preferred orientation *θ*_*k,OP*_ at cortical site *k* from the angle between the center-of-mass positions of ON/OFF RGCs.

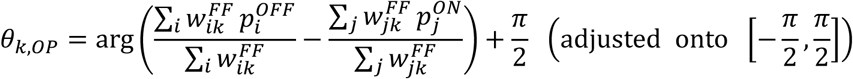

Then, we filtered the orientation preferences of cortical neurons using a 2D Gaussian filter of width *σ*_img_, obtaining an orientation map image *OP*(*x*) where *x* denotes pixel position. With the calculated orientation map and a given reference point *s*, we also computed an orientation similarity map as orientation preference similarity of all pixels *x* to *s*, given by

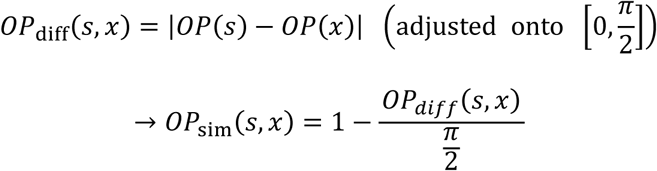

#### Simulation in salt-and-pepper organization

To show that patterned retinal waves can drive emergence of orientation-specific connections in salt-and-pepper organization of rodent V1, we performed additional modeling and developmental simulations of cortical a horizontal network using a mouse-data RGC mosaic^36^.

Following the notion of Jang et al. (2020)^54^, we modeled the rodent V1 by allocating sparse cortical sampling locations over the measured RGC mosaics of mouse^12,54,55^. Other than that, all the simulation settings were identical: the retinal waves were simulated over the RGC mosaic; the RGC-V1 feedforward wiring was further updated by retinal waves; and the horizontal connections were assigned and updated by retinal waves. For this model network, we only analyzed the orientation specificity of developed connections, because clustering index analysis and spontaneous activity simulation assumes the presence of a columnar orientation map. The simulation parameter details are provided in Table 1.

### Quantification and statistical analysis

#### Trend of developed V1 horizontal connection weights with respect to orientation

To assess how the newly developed V1 horizontal connections are related to the preferred orientations of the connected neurons, we first classified the weights of all nonzero horizontal connections into six groups according to the orientation difference between the connected neuron pairs, as described below.

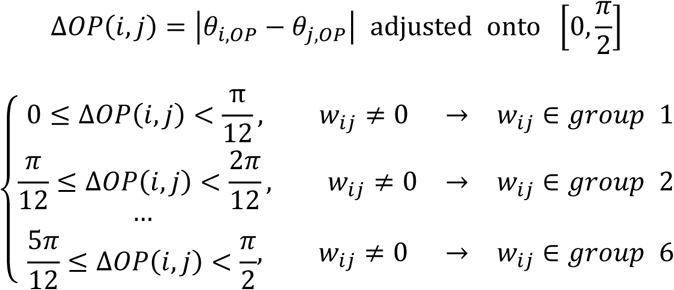

Then, we assessed the group trends using nonparametric Cuzick’s test for the trend of categorical data, under the null hypothesis that there was no trend across the groups. The p-value was calculated from the z-statistics given by the test. To rule out the effect of local connections, we did not consider connections between neurons within a distance of 200 µm (*i, j* such that 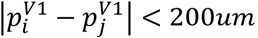) in our analysis.

For the mouse V1 model, we conducted an additional test for comparison with the experimental results of Ko et al. (2014)^37^. First, every cell pair (including pairs with zero connection weights) was categorized into one of three groups according to their orientation difference.

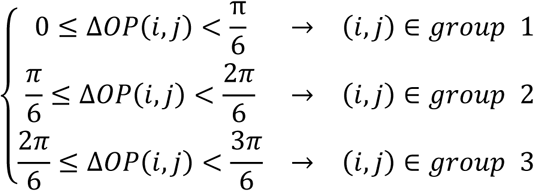

Then, we converted all the connection weight values to Boolean coupling relations by thresholding with 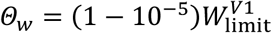. From that, we tested the trend of connection probability (# connected cell pairs in group / # cell pairs in group) across three groups using the Cochran-Armitage test for trend of categorical proportion, under the null hypothesis that there is no trend across the groups. The p-value was calculated from the z-statistics given by the test.

#### Spatial clustering of developed V1 horizontal connections

To quantify how much the developed cortical connections were spatially clustered in cortical domains, for a given V1 neuron, we selected 50% of the strongest-connected postsynaptic cortical locations. Then, for the selected locations, we sought to test whether the horizontal connection network was significantly clustered. Ruthazer et al. (1996)^3^ used Hopkins’ statistic with a sliding window to quantify clustering in the cell plots under the null hypothesis of spatial randomness ^3^. Here we summarize how we replicated their methodology.

For a set of points in a given window, two basic measurements were done: (1) A 10% random subset was taken from the point set, and nearest neighbor distances from each member in the random subset to the whole set were measured (denoted *w*). (2) A set of random locations (with set size same as in the first measure) within the window was selected, and nearest neighbor distances from the random locations to the whole point set were measured (denoted *x*). For the given window, a Hopkin’s statistic was computed as ln 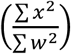.

Given a V1 neuron and 50% strongest postsynaptic locations in the cortical space, we moved a circular sliding window of radius 2*d*_*OFF*_ and collected H-statistic values at each sliding window position. Then, we took the median value of the H-statistics over every window as the clustering index (CI) of the presynaptic neuron’s cortical connectivity. We excluded the surrounding region of radius 2*d*_*OFF*_ from our analysis to check only for clustering in the remote postsynaptic area, considering the shape of the long-range connections in the V1 layer 2/3 of higher mammals.

For a developed V1 network, we obtained CI values from 50 randomly selected presynaptic neurons, following the above procedure. The same procedure was repeated for the initial random network and for the network developed from permuted retinal waves as well. Moreover, the significance of long-range clustering relative to the initial network was assessed using a two-sample t-test of CI values.

#### Matching between the spontaneous activity correlation map and orientation map

To quantify the alignment between the cortical activity correlation pattern and orientation map, for a given reference point *s*, we measured Pearson’s correlation coefficient between the orientation similarity map *OP*_*sim*_(*x*) and activity correlation map *C*(*x*) of reference *s* (*x* denotes pixel locations).

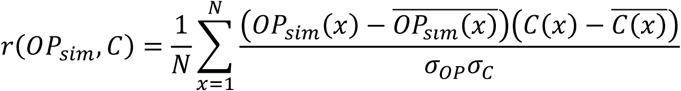

Here, *N* is the number of pixels. To test for statistical significance of the spatial correlation between the two maps, we made a control version of the correlation pattern *C*′(*x*) by randomly rotating the original *C*(*x*). Then, for the overlapping region *x*_*o*_ of rotated *C*′ and original *OP*_*sim*_, *r*(*OP*_*sim*_(*x*_*o*_), *C*(*x*_*o*_)) and *r* (*OP*_*sim*_(*x*_*o*_), *C′*(*x*_*o*_)) were measured. The same analysis was done 100 times with different control maps, and the values of *r*(*OP*_*sim*_, *C*) and *r* (*OP*_*sim*_, *C′*) were compared by paired t-test to generate p-value.

The above procedure assesses the significance of the correlation map and orientation map matching given a single, selected reference point. To test the global coherence of activity-orientation matching as well, we obtained *r*(*OP*_*sim*_, *C*) values from all locations of the V1 neurons as reference points and tested for significance using a two-sided t-test.

## Data and code availability

All the data supporting the findings of this study, model simulations, and data analysis codes are available on GitHub (https://github.com/jw9730/vsnn-retinal-wave).

## Key resources table

| REAGENT or RESOURCE | SOURCE | IDENTIFIER |
| --- | --- | --- |
| Software and Algorithms |  |  |
| MATLAB 2018a | MathWorks | RRID:SCR_001622 |

## Author contributions

**Jinwoo Kim:** Conceptualization, Methodology, Software, Writing, Visualization; **Min Song:** Conceptualization, Methodology, Software, Writing, Visualization; **Se-Bum Paik:** Conceptualization, Methodology, Writing, Supervision, Project administration, Funding acquisition

## Acknowledgments

This work was supported by the National Research Foundation of Korea (NRF) grants funded by the Korean government (MSIT) (No. NRF-2019R1A2C4069863, NRF-2019M3E5D2A01058328) (to S.P.).

## Supplementary information Videos

**Supplementary Video 1.** Sample retinal wave simulated on an extended mosaic of cats from Zhan et al. (2000)^50^.

**Supplementary Video 2.** Change of activity correlation pattern as reference point slides over cortical space. Simulation based on cell body mosaics in cats from Zhan et al. (2000)^50^.

**Supplementary Video 3.** Change of activity correlation pattern as reference point slides over cortical space. Simulation based on cell body mosaics in monkeys from Gauthier et al. (2009)^50^.

